# Phage-meropenem synergy against OXA-48-producing *Klebsiella pneumoniae* clinical isolates

**DOI:** 10.64898/2026.01.21.700822

**Authors:** Irene Cantallops, Celia Ferriol-González, Tamara Barcos-Rodríguez, Felipe Fernández-Cuenca, Javier E. Cañada-García, Silvia García-Cobos, Pilar Domingo-Calap

## Abstract

Antimicrobial resistance is a growing global healthcare crisis, driven by the rapid spread of resistant pathogens that compromise existing treatments. Carbapenem-resistant *Klebsiella pneumoniae* is a major public health threat, requiring novel therapeutic strategies. Phage therapy, which employs phages to target bacterial pathogens, is a promising approach, particularly when combined with antibiotics to enhance efficacy through synergistic interactions. In this study, time-kill curve assays were used to evaluate the synergy between the lytic phage vB_Kpn_2-P4 and meropenem against twelve *K. pneumoniae* clinical isolates from Spanish hospitals that carried diverse carbapenemases. Notably, in OXA-48-producing isolates, this combination prevented the emergence of resistant mutants, highlighting the therapeutic potential of phage-antibiotic synergy. The observed effect, linked to the presence of the pOXA-48 plasmid, suggests a promising strategy for combating multidrug-resistant bacteria.

## INTRODUCTION

Multidrug-resistant (MDR) bacteria pose a severe threat to public health, increasing morbidity, mortality, and healthcare costs (1). Among them, *Klebsiella pneumoniae* is a major threat due to its rapid spread and limited treatment options (2, 3). The World Health Organization (WHO) has designated carbapenem-resistant *K. pneumoniae* (CRKP) as a critical priority for new antibiotic development (4). Carbapenems, such as meropenem, have been a last-resort treatment for MDR Gram-negative infections. These antibiotics target bacterial penicillin-binding proteins, disrupting cell wall synthesis and leading to bacterial cell death (5). However, CRKP strains have evolved resistance through multiple mechanisms, including the production of carbapenemases such as KPC, NDM, VIM, and OXA-48 (6). These enzymes hydrolyze carbapenems and other β-lactams, often encoded on conjugative plasmids that facilitate rapid dissemination (7). The conserved pOXA-48 plasmid, carrying blaOXA-48, has been widely reported as a driver of resistance (8). The spread of high-risk CRKP clones, such as ST147, ST11, and ST15, has exacerbated the antimicrobial crisis, particularly in Southern Europe (9–11).

Given the stagnation in antibiotic development (12) and the increasing resistance burden, alternative therapies are urgently needed (13). One promising approach is phage therapy, which uses lytic phages to target bacterial pathogens (14). While once overshadowed by antibiotics, phage therapy has regained interest due to its specificity, adaptability, and potential to combat resistance (15, 16). Despite the effectiveness of phages against *K. pneumoniae in vitro* (17, 18), resistance can emerge rapidly. Combining phages with antibiotics has been shown to enhance bacterial clearance and reduce resistance development through phage-antibiotic synergy (PAS) (19, 20).

In this study, we evaluate the synergistic effects of meropenem and the broad host-range lytic phage vB_Kpn_2P-4 (18) against carbapenemase-producing clinical isolates. Our findings demonstrate PAS in OXA-48-producing strains, evidenced by bacterial load reduction and complete eradication at high antibiotic concentrations. These results highlight the potential of phage-antibiotic combinations as a strategy against MDR *K. pneumoniae*.

## RESULTS

### Variable Impact of Meropenem on clinical isolates

Clinical strains of *K. pneumoniae* with capsular type KL64, which is associated with high-risk clones (21) (Table 1), were grown in the presence of meropenem 20 and 100mg/L, respectively. Our results highlight variability in meropenem resistance (Figure 1). Some strains showed reduced OD_620_ values compared to the growth control, suggesting decreased viability at higher antibiotic concentrations. In contrast, others maintained or even increased OD_620_ values, indicating enhanced replication despite exposure to meropenem. Under our experimental conditions, all clinical isolates were able to grow at Css and, therefore, were considered meropenem-resistant according to our methodology. Minimum inhibitory concentration (MIC) values available on Table 1 may differ from resistant profiles in LB given the methodological variations.

**Figure 1.**
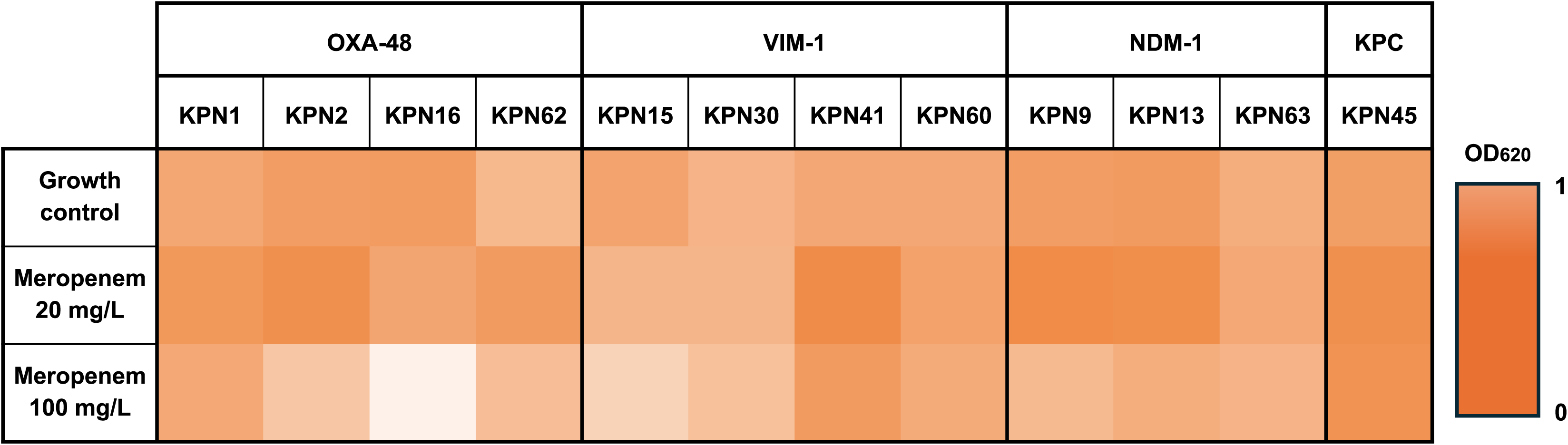
Effect of Meropenem (20 and 100 mg/L) on carbapenemase-producing clinical isolates, classified according to their carbapenemase type. Heat map represents the strength of lysis (OD_620_) measurement after 6 hours incubation at 37°C.

**Table 1.** Collection of clinical isolates of carbapenemase-producing *Klebsiella pneumoniae*.

|  | Isolate name | ST | Carbapenemase | ESBL/pAmpC | pAmpC | Narrow-spectrum beta-lactamase | Meropenem MIC (mg/L) | Biological origin | Source | Project | Accession ID |
| --- | --- | --- | --- | --- | --- | --- | --- | --- | --- | --- | --- |
| Kpn1 | 20180810 | 147 | OXA-48 | CTX-M-3 | - | - | ≤1 | Axillary smear | PIRASOA reference laboratory | PRJEB89930 | ERS24799066 |
| Kpn2 | 20180848 | 147 | OXA-48 | CTX-M-3 | - | OXA-1 | ≤1 | Urinary sample | PIRASOA reference laboratory | PRJEB89930 | ERS24799067 |
| Kpn9 | 20190469 | 147 | NDM-1 | CTX-M-15 | CMY-4 | - | 16 | Surgical device | Delgado-Valverde et al., 2023 | PRJEB53686 | ERR11279288 |
| Kpn13 | 20190525 | 147 | NDM-1 | CTX-M-15 | CMY-4 | - | 16 | Respiratory smear | Delgado-Valverde et al., 2023 | PRJEB53686 | ERR11281673 |
| Kpn15 | 20190643 | 15 | VIM-1 | CTX-M-15 | - | - | 2 | Unknown | Delgado-Valverde et al., 2023 | PRJEB53686 | ERR11281684 |
| Kpn16 | 20190646 | 147 | OXA-48 | CTX-M-15 | - | TEM-1 | 1 | Urinary sample | Ferriol-González et al., 2024 | PRJEB70314 | ERR12308420 |
| Kpn30 | 20200699 | 15 | VIM-1 | CTX-M-15 | - | TEM-1 | 2 | Urinary sample | Pérez-Palacios et al., 2023 | PRJEB52486 | ERR9741493 |
| Kpn41 | A1009KPN | 147 | VIM-1 | - | - | - | 2 | Urinary sample | Cañada-García et al., 2022 | PRJEB50822 | ERR8604201 |
| Kpn45 | A2608KPN | 11 | KPC-2 | CTX-M-65 | - | - | >16 | Blood sample | Cañada-García et al., 2022 | PRJEB50822 | ERR8604278 |
| Kpn60 | K11373 | 147 | VIM-1 | - | - | - | 16 | Rectal smear | Ferriol-González et al., 2024 | PRJEB68301 | ERR12251644 |
| Kpn62 | K11933 | 147 | OXA-48 | CTX-M-15 | - | - | >16 | Unknown | Ferriol-González et al., 2024 | PRJEB68301 | ERR12251646 |
| Kpn63 | K12100 | 147 | NDM-1 | CTX-M-15 | - | - | >16 | Unknown | Ferriol-González et al., 2024 | PRJEB68301 | ERR12251647 |

ESBL: Extended-Spectrum beta-lactamase; pAmpC: plasmid AmpC; MIC: minimum inhibitory concentration; -: Not present

### Lytic phage vB_Kpn_2-P4 efficiently infects high-risk clones

The lytic phage vB_Kpn_2-P4 was selected due to its broadest host range, infecting all the clinical isolates used in this study (18). Double-layer agar tests were performed to characterise the activity of the phage in solid medium for each strain, revealing different phenotypes. Despite slight variations in efficiency of plating (EOP), vB_Kpn_2-P4 exhibited lytic activity against 10 of the 12 clinical isolates on solid LB medium. Double-layer agar tests revealed diverse lysis phenotypes, with all OXA-48-producing strains showing complete lysis, while those carrying other carbapenemases displayed more variability. The turbidity of plaque halos suggests either partial bacterial lysis or phage-associated enzyme activity degrading bacterial cell wall components or the extracellular matrix. Growth curve analysis confirmed phage efficacy in liquid culture, demonstrating significant bacterial reduction in most strains (Figure 2). For example, Kpn30 exhibited over 50% growth reduction, whereas Kpn62 showed minimal impact. Notably, resistant mutant emergence varied, with a delay of nearly five hours between strains Kpn41 and Kpn63. These findings indicate that vB_Kpn_2-P4 effectively infects all clinical isolates in liquid culture, irrespective of their carbapenemase type.

**Figure 2.**
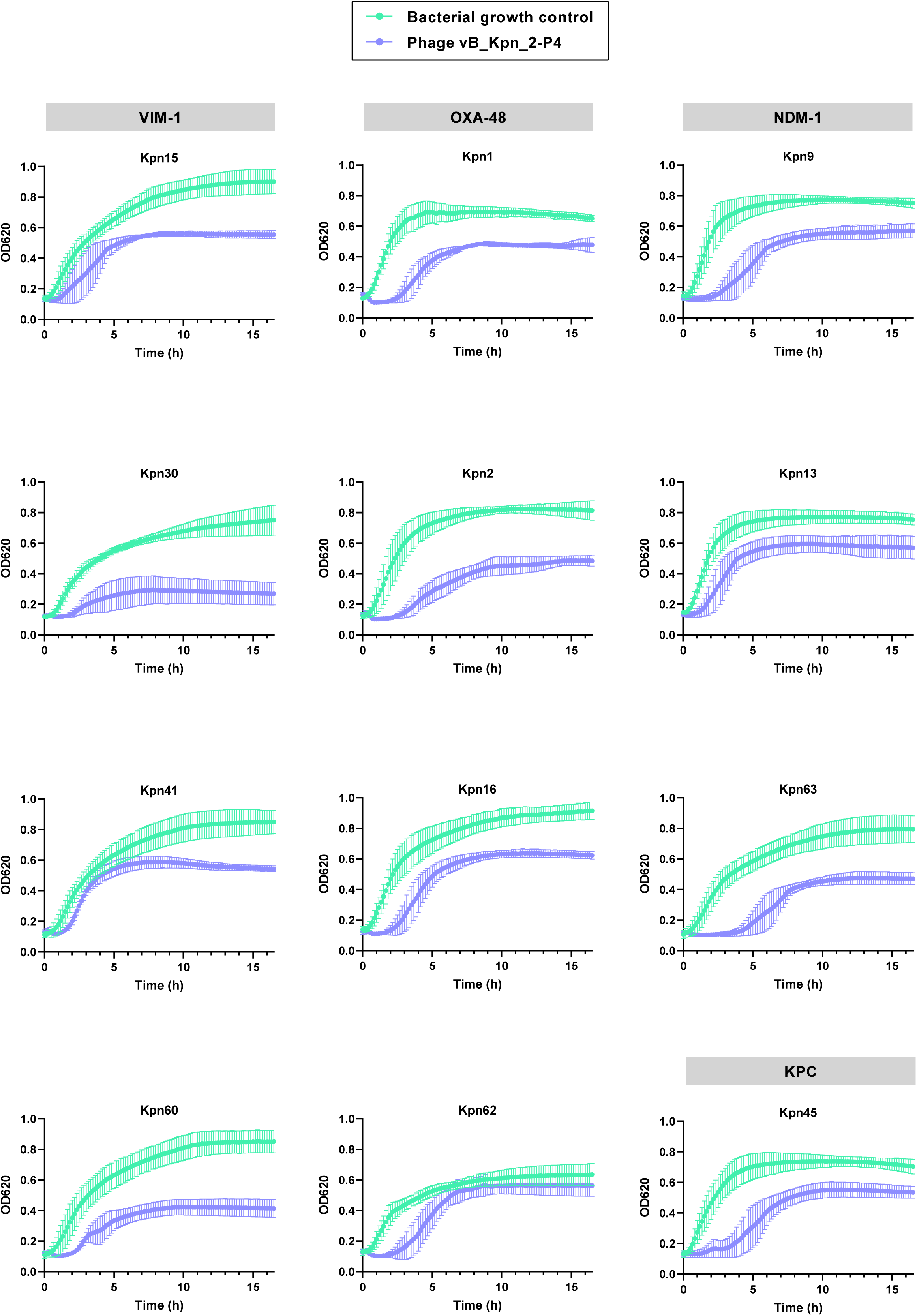
Bacterial growth over time assessed for 16.5 hours in the presence of phage vB_Kpn_2-P4 on clinical isolates harbouring different carbapenemases: OXA-48, NDM-1, VIM-1 and KPC. Growth curves show mean plus/minus standard deviations (SDs).

### Phage-antibiotic synergy between meropenem and vB_Kpn_2-P4 in clinical isolates carrying blaOXA-48 gene

To assess PAS, phage vB_Kpn_2-P4 was tested in combination with meropenem at concentrations of 20 mg/L and 100 mg/L. A significant PAS effect was observed exclusively in clinical isolates harbouring the *blaOXA-48 gene* (Figure 3), while no synergistic effects were detected in strains carrying other carbapenemases (Figure S1). Bacterial eradication was achieved when bacterial cultures were treated with 100 mg/L meropenem in combination with the phage, while 20 mg/L meropenem led to a substantial reduction in bacterial load, except for strain Kpn62. In this case, no significant difference was observed between phage treatment alone and its combination with 20 mg/L meropenem. However, at 100 mg/L, Kpn62 followed the same trend as the other OXA-48-producing strains, exhibiting complete bacterial clearance.

**Figure 3.**
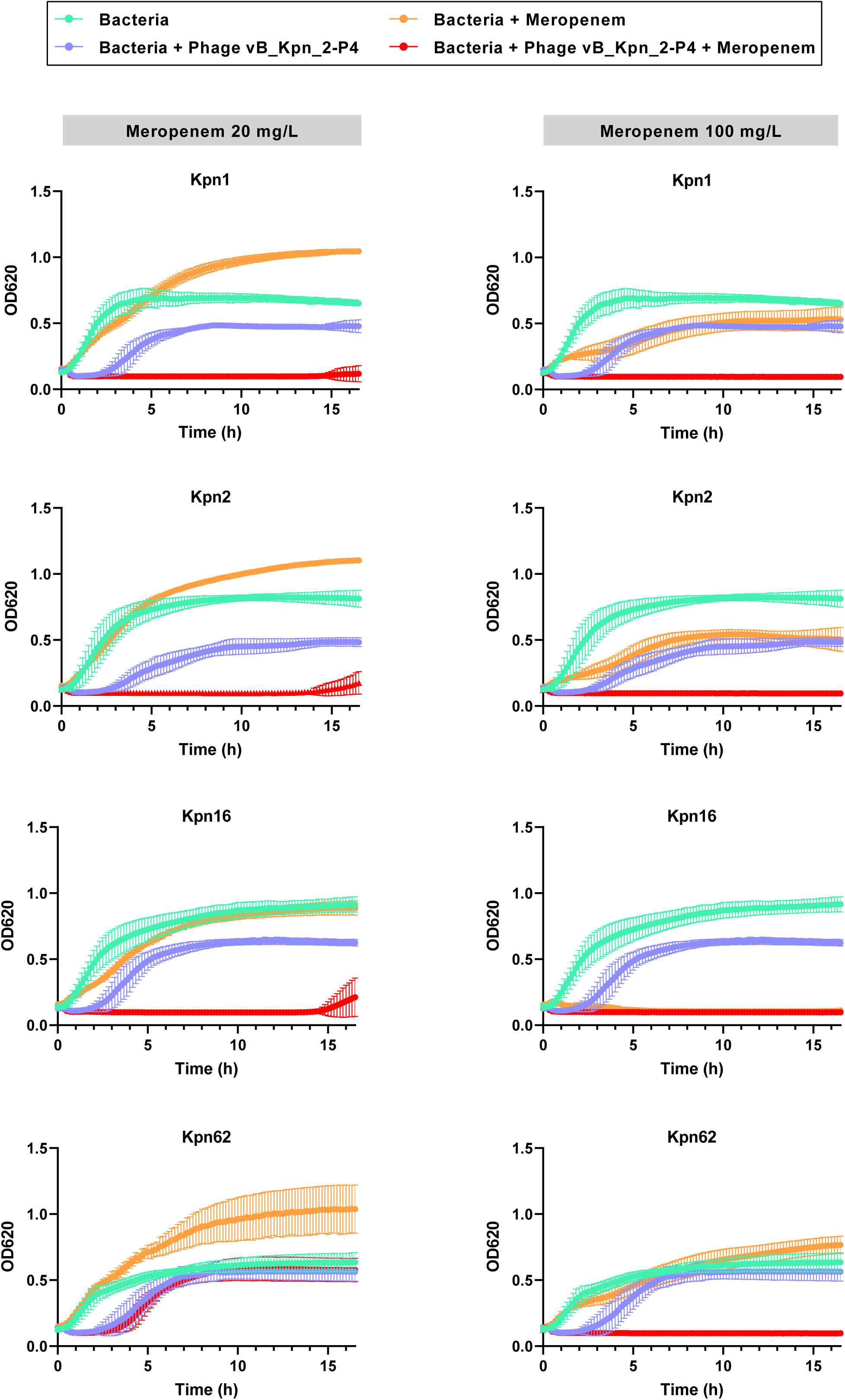
Bacterial growth over time assessed for 16.5 hours in the presence of phage vB_Kpn_2-P4 and/or meropenem 20 mg/L (left) and 100 mg/L (right) on clinical isolates harbouring OXA-48. Kpn1, Kpn2, Kpn16 and Kpn62. Growth curves show mean plus/minus standard deviations (SDs).

### Genomic basis of phage-antibiotic synergy in *blaOXA-48+* isolates

To investigate why the PAS effect was exclusive to *blaOXA-48+* strains, we performed a genome-wide association study (GWAS) on the bacterial isolates used in this study. Our analysis identified 74 CDSs associated with the PAS phenotype. Among them, one was a 46-amino acid hypothetical protein located on the chromosome, while the remaining 73 were encoded by the highly conserved conjugative plasmid pOXA-48 (Figure 4, Data S1 and S2). The limited number of samples, as well as the high level of conservation of this plasmid between the samples, might explain that practically the whole plasmid is associated with PAS in our GWAS. To determine whether PAS resulted from the loss of the *blaOXA-48* gene under selective pressure from phage vB_Kpn_2-P4, we performed PCR detection of the *blaOXA-48* gene in phage-resistant 10 colonies of each PAS-positive isolate: Kpn1, Kpn2, Kpn16, and Kpn62. Results confirmed that these resistant isolates retained the *blaOXA-48* gene, indicating that plasmid or gene loss was not responsible for the observed PAS effect. To verify the functionality of the *blaOXA-48* gene, the complete coding region was sequenced. No loss-of-function mutations were identified in any of the phage-resistant clones.

**Figure 4.**
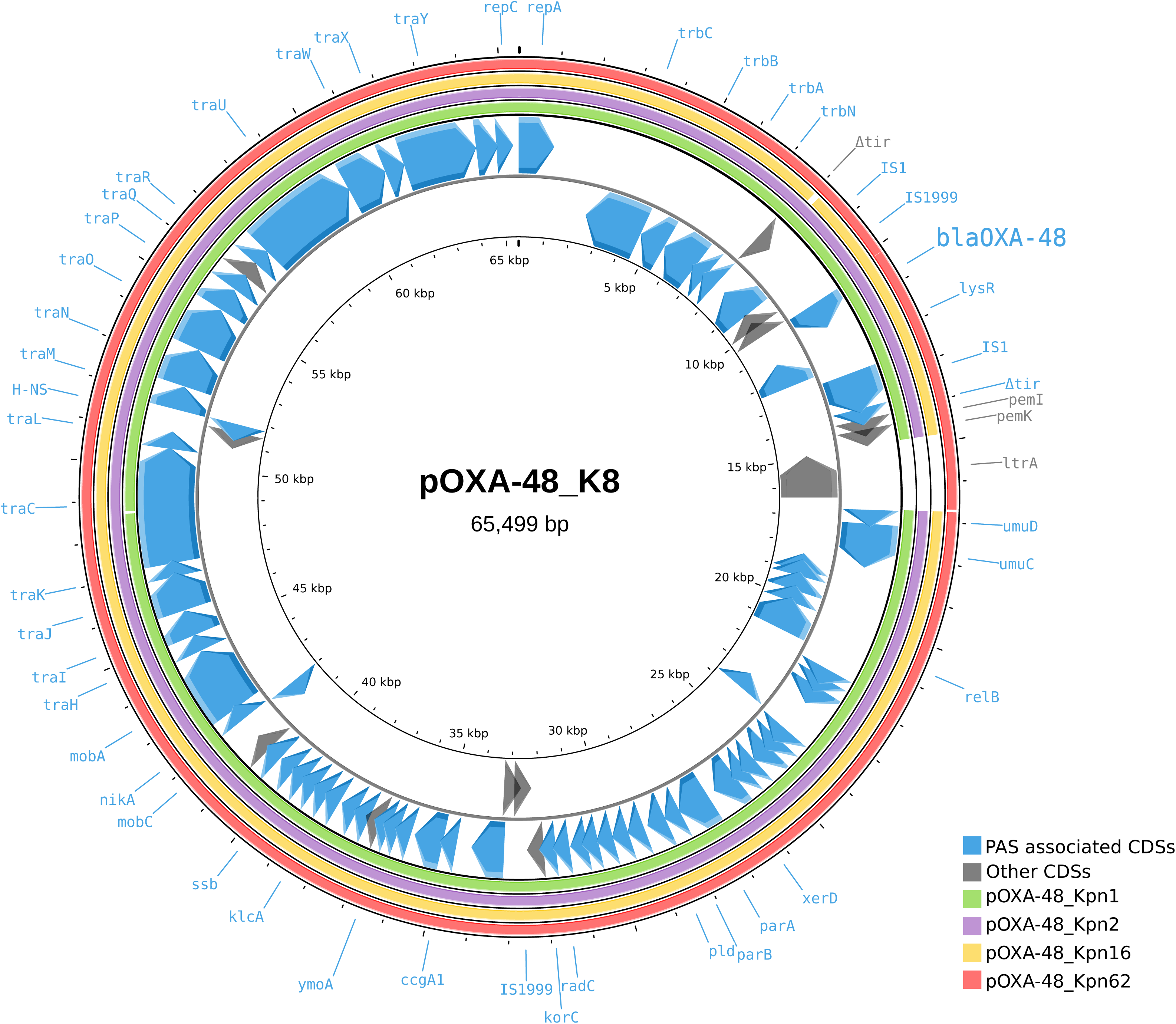
PAS-associated CDSs located in plasmid pOXA-48. Plasmid pOXA-48_K8 (MT441554.1) with its annotation by Toribio-Celestino et al., 2024 was employed as a reference. pOXA-48 contigs of sequenced OXA-48+ clinical isolates Kpn16 and Kpn62 were plotted against the reference and represented as external yellow and orange circles correspondingly. All genes associated with PAS by GWAS were indicated in blue.

## DISCUSSION

The emergence of extended-spectrum beta-lactamase and carbapenemase-producing *K. pneumoniae*, particularly those associated with the ST147 and KL64 capsular type, has led to hospital outbreaks worldwide (22–24). Notably, OXA-48 and its variants have become the dominant carbapenemases across Europe (25). The widespread dissemination of high-risk clones underscores the need for robust genomic surveillance and infection control strategies (10, 26). Carbapenem-resistant strains can survive even at the highest clinically permissible doses of meropenem. Paradoxically, antibiotic-induced stress can even promote bacterial growth in these resistant strains (27, 28). Our findings reveal that phage vB_Kpn_2-P4 can lyse ST147, ST15, and ST11 KL64 isolates, but PAS was observed exclusively in strains harbouring OXA-48.

Carbapenemases are primarily plasmid-borne, often imposing fitness costs on their bacterial hosts (7). The *blaOXA-48* gene, typically carried by the highly conserved conjugative plasmid pOXA-48, has been identified as a major contributor to these costs (29). The proposed mechanisms underlying this include envelope stress due to inefficient processing of the β-lactamase signal peptide (30) and structural alterations from DD-endopeptidase activity (31). These factors may select for the loss of the plasmid or the mutation of the β-lactamase producing gene in the absence of antibiotics, however, our results evidence that the bla-OXA-48 gene remains intact in phage-resistant clones of *blaOXA-48+* bacteria. Additionally, pOXA-48 harbours the insertion sequence IS1, which has been implicated in bacterial adaptation by inactivating the capsule operon, enhancing fitness under laboratory conditions (32). Notably, capsule loss is a common resistance mechanism against capsule-targeting phages (15), and our results suggest that both phage-resistant and phage-antibiotic-resistant colonies may be acapsular. However, these factors alone do not explain why PAS occurs exclusively in pOXA-48+ strains, highlighting the need for further investigation.

Summarizing, our study highlights the potential of phage vB_Kpn_2-P4 as an adjunct therapeutic agent against carbapenemase-producing *Klebsiella pneumoniae* in combination with meropenem. The observed PAS effect only in OXA-48-producing strains suggests a complex interplay between phage resistance mechanisms, plasmid carriage, and bacterial fitness costs. Our study provides key insights into the potential of combined phage-antibiotic therapy as an effective strategy against clinical isolates of OXA-48-producing *K. pneumoniae.* Future research should explore the efficacy of our phage and others from our collection in more clinical isolates. Additionally, testing the phage in combination with antibiotics with similar or distinct mechanisms of action could help elucidate the precise interactions between the phage and bacterial cells. This would advance our understanding of PAS and contribute to the broader application of phage therapy in treating resistant bacterial infections, particularly in clinical settings where conventional antibiotics are increasingly ineffective.

## METHODS

### Collection of *K. pneumoniae* clinical strains

Twelve clinical isolates of the KL64 capsular type were selected for this study (18). Detailed information on isolate designation, MIC, source, sample origin, sequence type (ST), beta-lactamase, and carbapenemase genes is provided in Table 1. All isolates were collected from clinical samples in Spanish hospitals. The high-risk clones ST147, ST15, and ST11, all associated with KL64, are highly prevalent in Spain and have acquired multiple carbapenemases (OXA-48, NDM-1, VIM-1, and KPC). The MICs were previously determined by the hospitals using the broth microdilution method in accordance with the recommendations of the European Committee on Antimicrobial Susceptibility Testing (EUCAST). Briefly, cation-adjusted Mueller–Hinton broth (CAMHB) was used, and bacterial suspensions were standardized to a final inoculum of 5 × 10⁵ CFU/mL. Microdilution plates were incubated at 35 ± 1 °C for 18 ± 2 h under aerobic conditions. For some bacterial isolates, a semi automatic method was employed with panels DKMGN (Sensititre). The MIC was defined as the lowest antibiotic concentration that completely inhibited visible bacterial growth. Quality control was performed using the EUCAST-recommended reference strains Escherichia coli ATCC 25922 and *K. pneumoniae* ATCC 700603. MIC values were interpreted according to the current EUCAST clinical breakpoint tables.

### Bacterial culture media and antibiotics

The strains were cultured in 3.5 mL of Luria-Bertani (LB) broth supplemented with 5 mM CaCl₂ and incubated overnight at 37°C with shaking at 180 rpm. Meropenem (purchased from Thermo Fisher) is a broad-spectrum β-lactam antibiotic commonly used to treat high-risk infections caused by MDR pathogens, including carbapenemase-producing *K. pneumoniae*. For this study, overnight bacterial cultures were diluted approximately 1:10 to achieve an initial optical density at 620nm (OD_620_) of 0.1–0.15, measured using a high-precision spectrophotometer (Implen^TM^). Meropenem was tested at concentrations ranging from 4 to 250 mg/L, with bacterial growth assessed after 24 hours of incubation. Experimental conditions were designed to mimic meropenem’s pharmacokinetic/pharmacodynamic (PK/PD) profile in human plasma, based on data from the Sociedad Española de Farmacia Hospitalaria (33). Specifically, our experiments replicated the peak plasma concentration (Cmax = 100 mg/L) and steady-state concentration (Css = 20 mg/L) observed after intravenous administration of 2 g every 8 hours.

### Lytic phage characterization

Phage vB_Kpn_2-P4 is a double-stranded DNA virus from the *Autographiviridae* family and *Przondovirus* genus (PP848852), originally isolated from sewage water (20). It was selected for this study due to its broad host range against the *K. pneumoniae* clinical strains included in this study. The phage encodes two depolymerase proteins that degrade the exopolysaccharidic capsule, facilitating bacterial infection. Additionally, anti-defense system analysis (34) identified an anti-restriction-modification system (35) and the gp4.5 anti-toxin-antitoxin system, which degrades the repressor of the antitoxin (36).

### Phage amplification and concentration

The seed aliquot of the phage vB_Kpn_2-P4 was amplified in the same bacterial strain in which it was isolated (Kpn2, 20180848) to obtain high-titer lysates (1.5 × 10^10^ PFU/mL). Based on previous data, the selected multiplicity of infection (MOI) was 10. The seed aliquot of the phage was amplified in LB media supplemented with CaCl_2_ in the same bacterial strain in which it was isolated to obtain high-titer lysates that were passed through 0.22µm pore filters to remove bacterial debris. Filtered lysates were concentrated to minimum volume using FluidPrep Concentrating Pipette Select with Ultrafilter Concentrating Pipette Tips and eluted in PBS FluidPrep Elution Fluid (Innova Prep, Drexel, MO) (37).

### Bacterial growth curves in the presence of meropenem and phage

Growth kinetics of *K. pneumoniae* isolates were assessed using a 96-well plate format, with incubation at 37°C inside a plate reader (Multiskan) for 16.5 hours. Bacterial lysis dynamics were determined through time-lapse turbidity measurements (OD_620_) recorded every 10 minutes. All experiments were performed in triplicate, and the resulting data were statistically analyzed using Python (version 3.8.10), with the matplotlib library (version 3.4.3) used for graphical representation and analysis. Each strain was tested under five experimental conditions: meropenem at 20 mg/L and 100 mg/L, phage alone, and phage in combination with meropenem at both concentrations. Positive and negative controls were included for comparison. As previously stated, the initial OD_620_ was adjusted to 0.1–0.15 (corresponding to approximately 1 × 10⁸ CFU/mL). The phage vB_Kpn_2-P4 was added at a MOI of 10, derived from an initial stock containing ∼1.5 × 10¹⁰ PFU/mL.

### Genome-Wide Association Study

The whole-genome sequence was available for eight of the bacterial isolates. FASTA files were annotated using Prokka (version 1.14.6) (38) and GFF3 annotation files were utilized for pan-genome analysis with Roary (version 3.12.0) (39). Isolate phenotypes were categorized based on the presence or absence of the PAS observed in the growth curves. To identify coding sequences (CDSs) associated with the PAS-positive or PAS-negative phenotype, Scoary (version 1.6.16) (40) was employed. Given the limited number of available genomes, a CDS was considered associated with a specific phenotype if it was present in all genomes exhibiting that phenotype (Naïve p < 0.05). Identified CDSs were further validated using blastp (BLAST 2.15.0+) (41). To determine whether selected proteins were encoded chromosomally or within mobile genetic elements, bacterial contigs were classified using Genomad (version 1.8.1) (42). The exact contig location of each selected CDS was confirmed using blast (41). Plasmidic contigs harboring selected CDSs were identified as components of the pOXA-48 plasmid based on sequence homology with *K. pneumoniae* strain K8 plasmid pOXA-48_K8 (MT441554.1), exhibiting a nucleotide sequence identity of 99.55–100%. For an accurate representation, the reference pOXA-48_K8 plasmid annotation was based on previous literature (7) and a blastp of all CDSs was performed to the PAS-associated CDSs.

### Sequencing of blaOXA-48 gene in phage-resistant bacterial colonies

An overnight culture of the PAS-positive isolates (Kpn1, Kpn2, Kpn16, and Kpn62) was grown in the presence of phage vB_Kpn_2-P4 in LB broth during 6h at 37 °C and 800 rpm. Phage-resistant bacteria were recovered by centrifugation and pellet resuspension. Phage-resistant phenotype was validated by serial dilution spot test of the recovered bacteria over a LB agar plate with a top agar monolayer of vB_Kpn_2-P4 phage. Phage-resistant bacteria were also plated in an LB agar plate for the recovery of single colonies. 10 phage-resistant and 3 wild type colonies were isolated. An 842 bp fragment including blaOXA-48 gene was amplified by PCR in all resistant and wild type isolated clones (Primers: >Forward CACACAAATACGCGCTAACCA, >Reverse GCATTAAGCAAGGGGACGTT) Phusion High-fidelity DNA polymerase (Thermo Scientific) was used for the reaction in a 25 µL final volume per sample (1 cycle of 98°C-30’’, 35 cycles of 98°C-10’’, 56°C 30’’and 72°C-55’’ and 1 cycle of 72°C-6’). Bands were detected by 1X agarose gel electrophoresis. All samples were positive, and amplicons were purified using the ZYMO DNA Clean and Concentrator-5 kit, and sequenced by Sanger sequencing.

## DATA AVAILABILITY

All data are presented in the manuscript and supplementary material.

## ACKNOWLEDGEMENTS

We thank the participating hospitals that submitted isolates to the Andalusian Reference Laboratory, Program for the Prevention and Control of Healthcare-Associated Infections and Antimicrobial Stewardship (PIRASOA, Servicio Andaluz de Salud) and the participating hospitals that submitted isolates to the Microbiological Surveillance Program on Antibiotic Resistance at the National Center of Microbiology. This research was funded by Agencia Estatal de Investigación and Ministerio de Ciencia e Innovación (project PID2020-112835RA-I00 and PID2023-150309OB-I00 funded by MCIN/AEI/10.13039/501100011033/). P. D-C was financially supported by a Ramón y Cajal contract RYC2019-028015-I (Spanish Ministry of Research and Innovation). C. F-G was funded by a PhD fellowship Atracció de Talent UV-INV_PREDOC-1913324 from Universitat de València.

## AUTHOR CONTRIBUTIONS

I.C. performed the experiments, data analysis, visualisation, and manuscript writing. C.F.-G. performed the computational analyses, data analysis, visualisation, and manuscript writing. T.B.-R. contributed to the experiments and visualisation. F.F.-C. contributed to bacterial resources, computational analyses, and manuscript revision. S.G.-C. contributed to bacterial resources and manuscript revision. J.E.C.-G. contributed to bacterial resources and manuscript revision. P.D.-C. provided reagents, conceived the project, designed the research, revised the manuscript, and conducted the supervision. All authors read and approved the final manuscript.

## COMPETING INTERESTS

P.D-C. is cofounder and scientific director of Evolving Therapeutics S.L. The other authors declare no competing interests.

