## Supplementary material for "Phage-meropenem synergy against OXA-48-producing *Klebsiella pneumoniae* clinical isolates": Figure S1

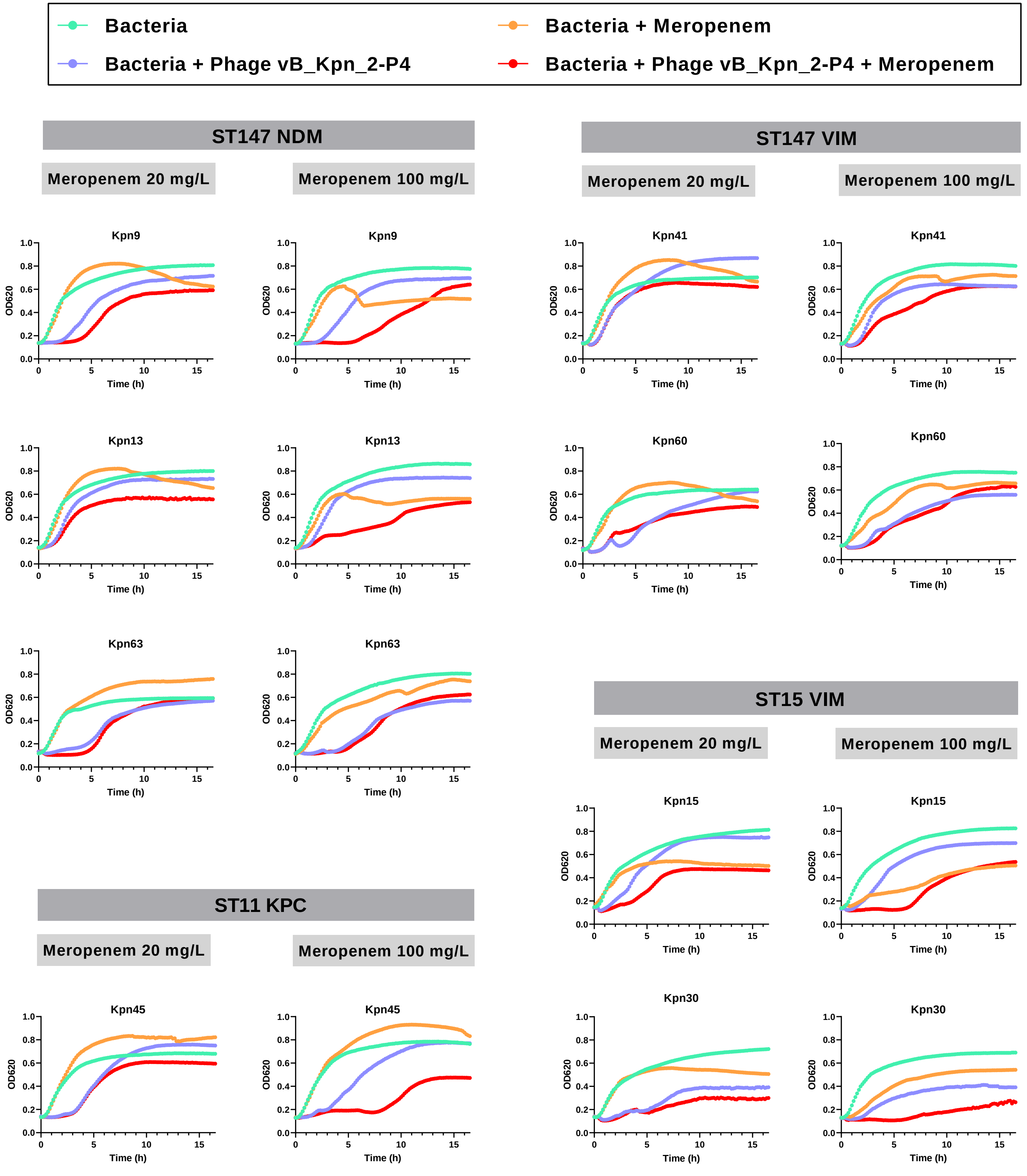


**Figure S1.** Bacterial growth (mean) over time assessed for 16.5 hours in the presence of phage vB_Kpn_2-P4 and/or meropenem 20 mg/L (left) and 100 mg/L (right) on clinical isolates harbouring other carbapenemases.
