## Supplementary material for "Phage-meropenem synergy against OXA-48-producing *Klebsiella pneumoniae* clinical isolates": Legends Data S1 and S2

**Supplementary Data**

**Data S1.** Bacterial gene association with phage-antibiotic synergy. PAS: Phage-antibiotic synergy.

**Data S2.** Gene presence-absence matrix including all gene groups built with the genome annotation of all isolates employed for the GWAS.
